# The molecular structure of *Schistosoma mansoni* PNP isoform 2 provides insights into the nucleotide selectivity of PNPs

**DOI:** 10.1101/300533

**Authors:** Juliana Roberta Torini, Larissa Romanello, Fernanda Aparecida Heleno Batista, Vitor Hugo Balasco Serrão, Muhammad Faheem, Ana Eliza Zeraik, Louise Bird, Joanne Nettleship, Yamini Reddivari, Ray Owens, Ricardo DeMarco, Júlio César Borges, José Brandão-Neto, Humberto D’Muniz Pereira

## Abstract

Purine nucleoside phosphorylases (PNPs) play an important role in the blood fluke parasite *Schistosoma mansoni* as a key enzyme of the purine salvage pathway. Here we present the structural and kinetic characterization of a new PNP isoform from *S. mansoni*, named as *Sm*PNP2. Screening of different ligands using a thermofluorescence approach indicated cytidine and cytosine as potential ligands. The binding of cytosine was confirmed by isothermal titration calorimetry, with a K_D_ of 27 μM, and kinetic parameters for cytidine catalysis were obtained by ITC resulting in a K_M_ of 76.3 μM. *Sm*PNP2 also displays catalytic activity against inosine and adenosine, making it the first described PNP with robust catalytic activity towards both pyrimidines and purines. Crystallographic structures of *Sm*PNP2 with different ligands were obtained and comparison of these structures with the previously described *S. mansoni* PNP (*Sm*PNP1) provided clues for the unique capability of *Sm*PNP2 to bind pyrimidines. When compared with the structure of *Sm*PNP1, substitutions in the vicinity of *Sm*PNP2 active site alter the architecture of the nucleoside base binding site allowing an alternative binding mode for nucleosides, with a 180° rotation from the canonical binding mode. The remarkable plasticity of this binding site deepens the understanding of the correlation between structure and nucleotide selectivity, offering new ways to analyses PNP activity.

**Author Summary:** *Schistosoma mansoni* is a human parasite dependent on purine salvage for purine bases supply. Purine nucleoside phosphorylase (PNP) is a key enzyme in this pathway. It carries two PNP isoforms, one previously characterized (*Sm*PNP1) and one unknown (*Sm*PNP2). Here we present the crystallographic structure of *Sm*PNP2 and its complex with cytosine, cytidine, ribose-1-phosphate, adenine, hypoxanthine, and tubercidin. Cytidine and cytosine were identified as ligands of *Sm*PNP2 using a thermofluorescence approach. Binding of cytosine was proven by Isothermal Titration Calorimetry (ITC) and cytidine, inosine, and adenosine kinetic parameters were also obtained. Purine bases showed different binding in the active site, rotated 180° from the canonical binding mode. It’s the first report showing a Low Molecular Mass PNP capable of catalyzing both types of nucleotide bases. The *Sm*PNP2 odd behavior sheds a new light on the *Schistosoma mansoni*’s life cycle metabolic adaptation.

## Introduction

Schistosomiasis is a tropical neglected disease that affects almost 206 million people worldwide according to World Health Organization (WHO), causing approximately 250,000 deaths annually (1). The disease is caused by a blood fluke of genus *Schistosoma* called *Schistosoma mansoni*, which is one of the main species being commonly found in Africa, Middle East, and the Americas.

It has been described that the *de novo* purine synthesis pathway is absent in adult and schistosomules life stages of *S. mansoni*, which depend exclusively on the purine salvage pathway (2–10). El Kouni and Naguib (11), investigated the pyrimidine salvage pathway “*in vivo*” and in extracts of *S. mansoni.* In intact worms, cytidine was the most efficiently incorporated pyrimidine precursor, while cytosine has failed to incorporate. The authors were unable to detect cytosine deamination to uracil and the cleavage of cytidine to cytosine.

Purine nucleoside phosphorylase (PNP) (E.C 2.4.2.1) is an enzyme responsible for reversible phosphorolysis of purine nucleosides, generating ribose-1-phosphate and their corresponding bases. PNPs belong to Nucleoside phosphorylase (NP-I) super-family and are classified into two groups: a Low Molecular Mass (LMM) group, composed of homotrimers with specificity for inosine, guanosine and their analogs and High Molecular Mass (HMM) group, composed of homohexamers with a broader specificity accepting both 6-oxo and 6-amino purines nucleosides and many others analogues (12). Other members of the NP-I family are the pyrimidine phosphorylases, for which Uridine Phosphorylase is the best-known example (13).

Recently Zhou et al. (14) have characterized three HMM NPs from thermophiles *Deinococcus geothermalis*, *Geobacilus thermoglucosidasius* and *Aeropyrum pernix.* All these NPs catalyze both 6-oxo and 6-amino purine substracts and some analogs, however, the natural substrates (inosine and adenosine) are better substrates than their 2-amino modified substrates. An intriguing discovery was the activity of these enzymes against cytidine, however, this activity (12-26 mU·mg^-1^) was several orders of magnitude lower than that observed for natural substrates (50-500 U·mg^-1^). The Nucleoside Phosphorylases can thus be thought as promiscuous enzymes involved in catalysis of one type of chemistry with many different substrates (15).

The structure and functional properties of the *S. mansoni* purine nucleoside phosphorylase isoform 1 (*Sm*PNP1) were reported by our group previously (16–24). The enzyme uses inosine and guanosine as substrates as it has been determined that the main determinant specificity of LMM PNPs towards 6-oxopurines is due to the presence of asparagine in the active site (N243 in human and N245 in *S. mansoni* PNPs). Site-direct mutagenesis, replacing the asparagine to aspartic acid in human PNP leads an increase of 5,000 fold of adenosine k_cat_ and a 4,000 fold increase of catalytic efficiency for adenosine (25).

Here, we characterize a newly identified isoform of *S. mansoni* PNP (*Sm*PNP2) employing a combination of methodologies: ligand discovery by thermofluorescence, kinetic assays, high-performance liquid chromatography (HPLC) and X-ray crystallography. Structural information of *Sm*PNP2 in both apo form and in complexes with cytidine, cytosine, ribose-1-phosphate, adenine, hypoxanthine, and tubercidin was obtained.

All these results allow us to describe for the first time the kinetics and structure of a LMM PNP well suited for inosine, adenosine and unexpectedly cytidine phosphorolysis. Finally, we also describe the structural basis for binding and catalysis of both purine and cytidine nucleosides.

## Material and Methods

### Cloning, expression, and purification of recombinant *Sm*PNP2

The *Sm*PNP2 gene (Smp_179110) was identified in the *S. mansoni* genome (26). The *Sm*PNP2 open reading frame (ORF) gene was synthesized with codon optimization by GenScript company and cloned into pOPINS3C (27) using the same protocol as previously described (28).

Two different protocols were employed for *Sm*PNP2 purification, one using OPPF_UK protocols, and the second utilized at Physics Institute of Sao Carlos (IFSC-USP), due to solubility problems with the OPPF protocol. In OPPF-UK, the cells were defrosted and lysed in lysis buffer (50 mM Tris-HCl pH 7.5, 500 mM NaCl, 30 mM imidazole and 0.2% Tween-20) supplemented with 25 μL/mL of cocktail protease inhibitors (Sigma-Aldrich) and 400 U/mL of DNAse I. Cells were lysed by Z cell disruptor at 30 kpsi and cell debris were removed by centrifugation (13,000 g for 45 min. at 4 °C). The clarified lysate was applied to a Ni^2+^-NTA column (GE, Healthcare) connected to an AKTA-Xpress purification system. Recombinant 6His+SUMO-*Sm*PNP2 was eluted with elution buffer (50 mM Tris pH 7.5, 500 mM NaCl, 500 mM imidazole) and injected into a size exclusion S200 column, pre-equilibrated with Gel filtration buffer (20 mM Tris pH 7.5, 200 mM NaCl and 1 mM Tris (2-carboxyethyl)phosphine (TCEP)). Fractions containing 6His+SUMO-*Sm*PNP2 were pooled and concentrated to 2-3 mg/mL.

In the second purification protocol, the cells were defrosted and lysed in a different Lysis buffer (50 mM potassium phosphate pH 7.4, 300 mM NaCl, 10 mM imidazole, 5 mM β mercaptoethanol, 0.1 mM dithiothreitol (DTT) and 1 mM phenylmethylsulfonyl fluoride (PMSF)). Cells were lysed by sonication followed by centrifugation (1,300 g for 45 min. at 4 °C). The clarified lysate was applied to Co-NTA agarose column (Qiagen), after column wash, the 6His+SUMO-*Sm*PNP2 was step-eluted with elution buffer (50 mM potassium phosphate pH 7.4, 300 mM NaCl, 200 mM imidazole and 5 mM β mercaptoethanol). Purified protein was visualized on a 15% SDS-PAGE after both purifications.

In both of the purification protocols, the 6His-SUMO tag was cleaved by addition of 3C protease (1 U/100 μg fusion protein) with overnight incubation at 4 °C. Cleaved *Sm*PNP2 was purified from the digest using a reverse purification in Ni^2+^-NTA column, the collected eluate was then concentrated and followed by dialysis in Tris-HCl buffer (20 mM Tris pH 7.4, 200 mM NaCl and 1 mM TCEP). Purified protein was concentrated up to 5-6 mg/mL in the appropriate buffer for further studies. All stages of *Sm*PNP2 production were visualized on a 12% SDS-PAGE.

### Differential scanning fluorimetry (DSF)

Employing the same procedures for activity assay used for *Sm*PNP1 (18), no activity for inosine phosphorolysis was observed for *Sm*PNP2. For this reason, DSF experiments were employed in order to identify the possible ligand for *Sm*PNP2.

Firstly, optimal concentrations of *Sm*PNP2 and SYPRO orange dye were determined to perform DSF on a grid of varying concentrations of *Sm*PNP2 protein and SYPRO orange in the Analysis Buffer (20 mM potassium phosphate pH 7.4, 300 mM NaCl. BioRad CFX96 thermocycler was then programmed to equilibrate sample at 25 °C for 5 minutes followed by an increase in temperature up to 95 °C, at a rate of 0.5 °C/min. The best condition found was 6 μM of *Sm*PNP2 and SYPRO orange 1:2000 dilution (dye: analysis buffer).

For ligand finding the Silver Bullets Bio kit (Hampton Research) was used. The reaction was performed using the best condition obtained as described above and contains 2 μL of Silver Bullets bio, in a final volume 20 μL. Apparent melting temperature (T_m_) was calculated from the mean of triplicate measurements and, the ΔT_m_ was calculated based on the difference of the T_m_ value from controls in absence of ligands and, the T_m_ value from the samples employing the DSF Melting Analysis (DMAN) software (29).

### Isothermal Titration Calorimetry

All of the ITC binding and kinetics experiments were carried out with an iTC200 microcalorimeter (GE Healthcare Lifesciences). Protein, ligands, and/or substrates were prepared in 50 mM potassium phosphate buffer (pH 7.6) plus 300 mM NaCl.

### Binding Experiments

In order to verify the potential of the cytosine base as a substrate for *Sm*PNP2, a preliminary binding experiment was performed. The sample cell was filled with 60 μM of protein (in the monomer concentration) and it was titrated with multiple injections of 1 mM cytosine, at 25 °C. The enthalpy for each injection was calculated by integrating the area under the peaks obtained from the change of power as a function of time. The heat of the injectant dilution was determined from the end of the titration and was subtracted from the data. The experimental isotherms were analyzed to yield apparent association constant (K_A_), apparent enthalpy change (ΔH_app_) and stoichiometric coefficient (n). The apparent Gibbs energy change (ΔG_app_) was calculated using the relation Δ*G_app_* = –*RTlnK_A_* where R is the gas constant (cal.mol^-1^.K^-1^) and T the absolute temperature (K). The apparent entropy change (ΔS_app_) was estimated by the follow equation Δ*G_app_* = Δ*H_app_* – *T*Δ*S_app_*. The apparent dissociation constant (K_D_) was calculated as the inverse of the K_A_ value.

### Kinetic Experiments

Two different experiments were performed to determine the kinetic parameters of the cytidine nucleoside phosphorolysis by *Sm*PNP2. For determination of the apparent enthalpy change ΔH_app_ for the phosphorolysis, 13 μM *Sm*PNP2 was titrated with 3 injections of 5 μL cytidine nucleoside 5 mM, at 20 °C where the substrate was quickly and completely converted into product. The integral of the area under each peak yields the ΔH_app_ of the reaction. For determination of the rate of substrate reaction, 500 nM *Sm*PNP2 was titrated with 25 injections of 1 μL cytidine 5 mM, at 20 °C. After the correction for the heat of the titrant dilution, these data were analyzed using the Origin 7.0 to obtain values for K_M_, V_max_ and k_cat_ for the enzymatic reaction, as indicated by the supplier. The kinetic parameters were determined as the average values of the parameters obtained after analysis of 3 experiments.

### *Sm*PNP2 HPLC data analysis

The HPLC was used to identify the *Sm*PNP2 catalytic reaction from cytidine to cytosine in the presence and absence of phosphate. Firstly, the reaction was prepared in 50 mM potassium phosphate buffer pH 7.4 with 1 mM cytidine and started with the addition of 200 nM *Sm*PNP2 at room temperature. Aliquots were withdrawn from time 0 to 40 minutes of the reaction (T_1_, T_2_, T_5_, T_10_, T_15_, T_20_, T_30_ up to T_40_, respectively), frozen in liquid nitrogen and heating at 85 °C for 15 minutes to denature the *Sm*PNP2 and followed to centrifugation at 10000 rpm for 20 minutes at 4 °C. The controls: phosphate buffer; cytidine; cytosine; ribose 1-phosphate (R1P) and a mix of all these substrates were prepared in the same buffer and submitted to same procedures.

An aliquot of 10 μL was applied in Supelcosil^™^ LC-18-S (Sigma-Aldrich) column coupled to HPLC Water system with 1 mL/min. flow and isocratic gradient using 88% Buffer A (97.5 mM potassium phosphate buffer pH 4.0 and 2.5 % methanol) and 12% Buffer B (97.5 mM potassium phosphate buffer pH 4.0 and 25 % methanol) monitored at 253 nm.

The reverse reaction using 1 mM cytosine in presence of 1 mM R1P was prepared in 50 mM HEPES buffer pH 7.4 at room temperature. The enzymatic reaction was started with 200 mM *Sm*PNP2 and monitored likewise previous mentioned and injected in the Supelcosil^™^ LC-18-S column to it analyze procedure by HPLC.

### Adenosine and inosine phosphorolysis assays

Using an enzyme amount three-fold higher than previously described for *Sm*PNP1 assays (18), we were able to detect kinetic parameters for adenosine and inosine. The kinetic parameters for adenosine were measured via coupled assay by xanthine oxidase. In this method, the xanthine oxidase converts free adenine from adenosine (Ado) to 2,8-dihydroxyadenine, resulting in an increase in the absorbance at A_305_ nm (ε = 15,500 AU) (30, 31). The kinetic parameters for inosine were also measured using coupled assay by xanthine oxidase, which converts the hypoxanthine formed into acid uric, resulting in an increase in the absorbance at A_295_ nm (ε = 14,000 AU).

Kinetic parameters were calculated in sextuplicate (from three independent experiments) at room temperature in a 200 μL reaction mix containing 100 mM potassium phosphate buffer at pH 7.4, twelve concentrations of substrate (4.5 to 400 μM) and 0.3 units of xanthine oxidase from bovine milk (Sigma-Aldrich). The reaction was started by adding 1 μM of *Sm*PNP2 to the reaction mixture, and the OD was immediately monitored using a SPECTRAmax^®^ PLUS384 spectrophotometer (Molecular Devices, USA). The kinetic parameters (K_M_ and k_cat_) were derived from non-linear least-squares fits of the Michaelis-Menten equation in the GraphPad Prism software using the experimental data.

### Crystallization, data collection, structure solution and refinement

Initially, a Pre-Crystallization Test (PCT) was performed (using *Sm*PNP2 in Tris buffer pH 7.5), to determine the optimal concentration of *Sm*PNP2 for setting up crystallization experiments. Using *Sm*PNP2 in concentration 5 mg/mL, crystallization screening experiments were performed on Cartesian 2 (OPPF-UK) using screen solutions from Molecular Dimension (Morpheus^®^ and JCSG-*plus*^TM^), Hampton Research (Index HT^TM^) and Jena Bioscience (JBScreen Wizard 1 and JBScreen Wizard 2) in Greiner Crystal-Quick 96 well-sitting drop plates (Hampton research, USA). Plates were incubated at 20 °C in FORMULATRIX imager (USA).

For co-crystallization experiments the *Sm*PNP2 at 5 mg/mL was incubated with 5 mM of each ligand (R1P, inosine, hypoxanthine, cytidine, and cytosine), and submitted to crystallization trials using Morpheus and JCSG-*plus* with 1:1 μL drops in Honeybee 961 robot in Greiner CrystalQuick plates and incubated at 18 °C. In all cases, cubic shape crystals appeared after two days and reached their maximum size (~150 μm) in three days.

Crystals used in soaking experiments were grown in 100 mM Bis-Tris pH 6.5, 200 mM ammonium acetate and 30% PEG3350. Soaking was carried out in mother solution (2 μL) plus 0.5 μL of ligand (30 mM of cytidine, 6 mM of hypoxanthine, 50 mM of tubercidin and 6 mM of adenine) dissolved in 100% DMSO, for one 1 hour. The crystals were mounted in micro loops, cryo-protected with 20% glycerol or PEG200 (if necessary) and cooled in liquid nitrogen. Diffraction data were collected using synchrotron radiation at beamlines I04 and I04-1 at Diamond Light Source (DLS, Harwell, UK). These data were indexed, integrated and scaled using Xia2 (32–35).

The *Sm*PNP2-apoenzyme structure was solved by molecular replacement using the Phaser program (36), employing *Sm*PNP1 isoform (PDB ID 3FB1) (24) which shares 61% identity as a search model. The remaining structures were also solved by molecular replacement, using one of the structures previously refined as a search model. The refinement was carried out using Phenix (37) and the model building was performed with COOT (38), using a weighted 2Fo–Fc and Fo–Fc electron density maps. In all cases the behavior of R and R_free_ was used as the principal criterion for validating the refinement protocol and the stereochemical quality of the model was evaluated with Molprobity (39). The coordinates and structure factors have been deposited into the Protein Data Bank (PDB) under the following codes: *Sm*PNP2 apo – 5CXQ, *Sm*PNP2 MES complex - 5CXS, *Sm*PNP2 cytosine complex – 5KO5, *Sm*PNP2 cytosine and R1P complex −5KO6, *Sm*PNP2 tubercidin complex – 5TBV, *Sm*PNP2 adenine complex – 5TBS, *Sm*PNP2 cytidine complex - 5TBT and *Sm*PNP2 hypoxanthine complex – 5TBU.

### Whole-mount *in situ* hybridization (WISH)

To produce the antisense RNA probes, DNA templates corresponding to segments of 200-300 bp of *Sm*PNP1 (Smp_090520) and *Sm*PNP2 (Smp_179110) were amplified from adult worms cDNA, using the following primers: PNP1 forward 5’-TGTCGAAAGCGATTTGAAGC-3’ and reverse 5’-TCATTTCAGCAAGTACACAAAGAGA-3’ and PNP2 forward 5’-CCATGAAATAGTTACTCGTTCTAACAA-3’ and reverse 5’-TGTGACCCGAAAAATTTGTAATG-3’. The primers were designed to comprise of untranslated regions (UTR) in order to avoid cross-reaction of the probe due to the sequence similarity. The amplicons were cloned into the pGEM-T-Easy vector (Promega), which contains both T7 and SP6 promoters; the orientation of the strands was verified by DNA sequencing. Transcription reactions to synthesize digoxigenin (DIG)-labelled RNA probes were performed using the Riboprobe kit (Promega) and the clones in pGEM-T as templates. The WISH protocol was carried out according to *in situ* hybridization optimized for *S. mansoni* (40, 41). Briefly, formaldehyde fixed adult worms were partially digested with proteinase K, incubated overnight at 56 °C in hybridization buffer containing the DIG-RNA probes (1 μg/mL). After being extensively washed, the worms were blocked with 10% horse serum in MABT (100 mM maleic acid, 150 mM NaCl, 0.1% Tween-20, pH 7.5), incubated overnight at 4 °C with anti-digoxigenin-AP Fab fragments (Roche) at 1:2000 dilution and revealed with NBT/ BCIP solution (Roche). The control group was incubated in hybridization buffer with sense DIG-RNA probe. Images were acquired using an Olympus BX53 microscope.

## Results and Discussion

### *Sm*PNP2 sequence and expression profile are markedly distinct from that of *Sm*PNP1

The sequence of *Sm*PNP2 was found searching the *S. mansoni* genome using GeneDB tools (42). The sequence of *Sm*PNP2 encodes for a protein with 287 amino acids (the same size as *Sm*PNP1 protein), with a calculated molecular weight of ~31.3 kDa. The identity between the protein sequences of the two *S. mansoni* PNPs is 61% (Supplementary Figure 1) and *Sm*PNP2 has 46% identical residues to its human counterpart.

Comparison of expression profiles derived from previous RNA-Seq experiments (26) available at GeneDB showed that *Sm*PNP1 and *Sm*PNP2 presented a different expression pattern: *Sm*PNP1 was highly expressed in adults while *Sm*PNP2 is more abundant in cercariae (with 24x more transcripts than an adult worm). Moreover, WISH experiments using a probe specific for *Sm*PNP1 in adult worms resulted in a strong staining of female vitelaria (Supplementary figure 2), indicating a specific role of this protein in the parasite reproduction and justifying its enrichment in adult worms. In contrast, WISH experiments using *Sm*PNP2 probes in adult worms were not successful, thus suggesting a low abundance of transcripts in this life stage. These data point out that *Sm*PNP1 and *Sm*PNP2 must be acting in very different and specific contexts and that these proteins may present very distinct characteristics to fulfill their roles. Considering this scenario, a thorough investigation of *Sm*PNP2 proprieties was considered to be desirable to have further information about its possible role.

### *Sm*PNP2 is able to bind cytosine

The *Sm*PNP2 gene was synthesized with codon optimization and cloned in five pOPIN vectors (43). After expression screening using several pOPIN vectors, we found that vector pOPIN-S3C, producing a fusion protein with SUMO, displayed best yields. After cleavage with PreScission protease, the *Sm*PNP2 was affinity purified using cobalt column. The yield was approximate 4 mg/L of culture medium.

Preliminary assays using *Sm*PNP2 showed very low activity against inosine (19 times lower than for *Sm*PNP1), which is a natural substrate for LMM PNPs. This unexpected characteristic of *Sm*PNP2 encouraged us to perform a DSF screening incubating the enzyme with mixtures containing several different compounds in order to find possible ligands.

An increase of T_m_ temperature was obtained in eight conditions varying from 2.5 to 0.5 °C. Analyzing the composition of these mixtures that led to an increase in melting temperatures, we observed that with exception of one mixture (composed by L-carnitine, tannic acid, aspartame, caffeine, p-coumaric acid, 4-hydroxyl-proline and Hepes), all the remaining conditions contained cytidine or cytosine. Curiously, no significant T_m_ increase was observed when *Sm*PNP2 was incubated with mixtures containing purines, suggesting a more unstable binding with such compounds.

The low of activity against inosine (in comparison to *Sm*PNP1 (19)) and the possible binding with cytidine and cytosine were completely unexpected. To our knowledge, no LMM PNP have been shown to have activity against pyrimidine bases or their nucleosides. Furthermore, in the canonical metabolic pathways cytidine is never directly converted to cytosine. Instead, it is either converted directly to CMP by uridine-cytidine kinase (E.C 2.7.1.48) or is deaminated to uridine by cytidine deaminase (E.C 3.5.4.5).

Isothermal Titration Calorimetry using cytosine and *Sm*PNP2 confirmed binding displaying a stoichiometric coefficient of 1 base per protomer and a K_D_ equal to 27 (± 3) μM, the interaction being both enthalpic and entropy (Table 1 and Supplementary Figure 3).

**Table 1.** Thermodynamic parameters of the interaction between S*m*PNP2 and cytosine, obtained by Isothermal Titration Calorimetry

| Parameter | Value |
| --- | --- |
| Binding Sites (n) | $0.71 \pm 0.07$ |
| $\Delta H_{app}$ (cal/mol) | $-4700 \pm 100$ |
| $\Delta S_{app}$ (cal/mol/deg) | $+5.14 \pm 0.06$ |
| $K_A$ ($M^{-1}$) | $37000 \pm 4000$ |
| $K_D$ ($\mu M$) | $27 \pm 3.0$ |

### *Sm*PNP2 efficiently catalyzes the phosphorolysis of cytidine

Considering the verified affinity of *Sm*PNP2 to cytosine, an obvious question is whether this protein has the capacity to catalyze the phosphorolysis of cytidine into cytosine. To address that, an HPLC analysis of a cytidine solution incubated with *Sm*PNP2 was performed running aliquots corresponding to different times of incubation and monitoring the column eluate at 253 nm to detect nucleotides. Comparison of retention times for cytidine and cytosine were determined using standards and allowed to deduce an increase of cytosine concomitantly to a decrease in cytidine absorbance during the experiment time (Figure 1A). The reverse reaction was also prepared and analyzed using the same procedure and indicated a reduced efficiency in cytidine formation from cytosine in comparison with the rate observed for cytidine phosphorolysis (Figure 1B).

**Figure 1.**
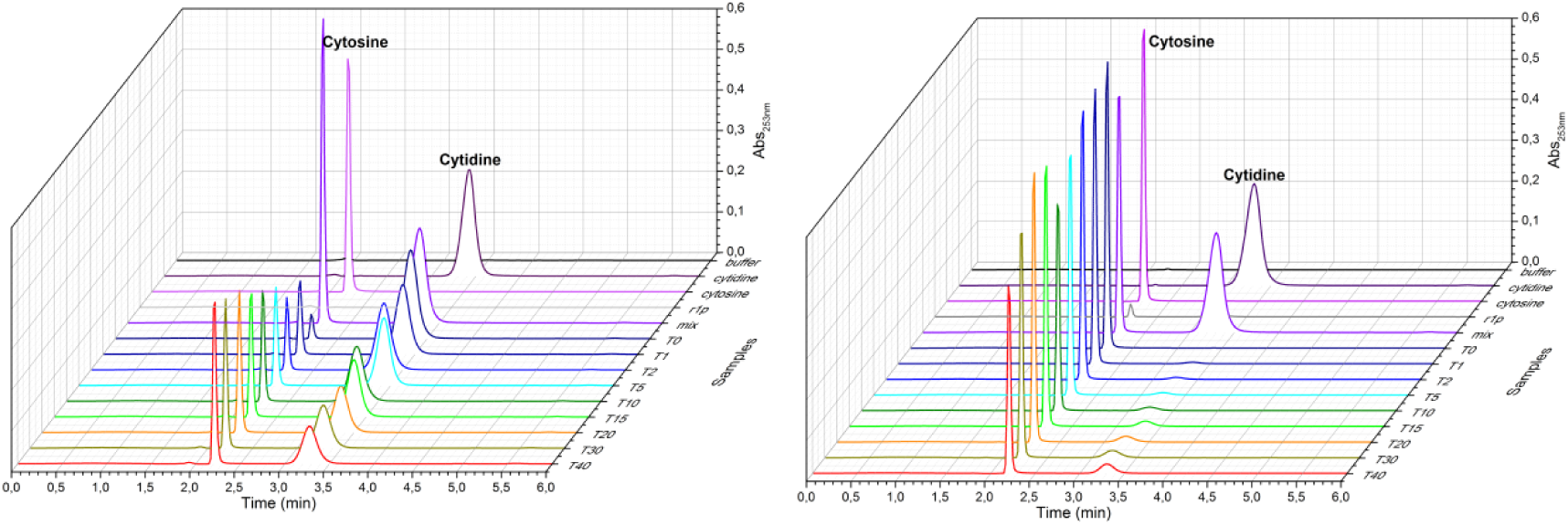
The HPLC chromatogram. **A.** The cytosine formation from cytidine in PPi presence, highlighting the cytosine formation (absorbance at 253 nm increase at 2.3 min). **B.** The reverse reaction in HEPES buffer showing the inefficiency of *Sm*PNP2 to this enzymatic reaction by the conservation of absorbance that represents the cytosine.

In addition, preliminary ITC binding assays of *Sm*PNP2 with cytidine showed that was not possible to achieve saturation even after performing assays in several experimental conditions. Indeed, analysis of the titration showed a profile compatible with the catalysis of the titrator, thus providing further direct evidence of cytidine nucleoside phosphorolysis by *Sm*PNP2. The kinetics of enzyme-catalyzed reaction was obtained through two independent calorimetric experiments. In the first experiment, the ΔH_app_ was determined with high concentrations of enzyme in the cell, relatively low amounts of cytidine in the syringe and allowing sufficient time between the injections to ensure that all of the substrates were converted to product (Supplementary Figure 4, *inset*). The obtained peaks were used to deduc the ΔH_app_ value for the catalytic reaction. The same experiment performed for purines (adenosine, guanosine, and inosine) and pyrimidines (thymidine and uridine) nucleosides did not evolve heat indicating the absence of detectable phosphorolysis in the tested conditions (data not shown). In the second experiment, the rate data was obtained with low amounts of *Sm*PNP2 in the cell and high amount of cytidine nucleoside in the syringe, ensuring that steady-state conditions are maintained while substrate concentration is kept almost constant (Supplementary Figure 4A). The data reaction obtained through the titration was fitted with the Michaelis–Menten model, allowing the calculation of the kinetic parameters (Table 2 and Supplementary Figure 4B). The K_M_ value obtained for the cytidine phosphorolysis (76.3 ± 0.3 μM) presents an intermediate value in comparison with spectrophotometric assays for inosine (136 ± 7 μM) and adenosine (22 ± 2 μM), respectively. In the same way, parameters related to the catalytic efficiency as k_cat_ and k_cat_/K_M_ are also similar for cytidine, inosine and adenosine phosphorolysis (Table 2).

**Table 2.** Kinetic parameters of the phosphorolysis promoted by S*m*PNP2.

| Substrate | $K_M$ ($\mu M$) | $k_{cat}$ ($s^{-1}$) | $V_{max}$ (mM/s) | $k_{cat}/K_M$ ($mM^{-1}.s^{-1}$) |
| --- | --- | --- | --- | --- |
| Adenosine | $22.3 \pm 2.5$ | $1.02 \pm 0.02$ | 0.001 | $46.0 \pm 5.2$ |
| Inosine | $136.4 \pm 7.0$ | $7.0 \pm 0.2$ | 0.006 | $50.6 \pm 2.9$ |
| Cytidine | $76.3 \pm 0.3$ | $2.0 \pm 0.2$ | 0.001 | $26.2 \pm 0.1$ |

It should be noted the only PNPs previously described to be capable of cytidine phosphorolysis displayed an activity several orders of magnitude lower than that observed for purines. In contrast, *Sm*PNP2 did not display a strong preference for purines in relation to cytidine and the relative concentration of nucleotides seems to be the determining factor. Therefore, it is possible argue that *Sm*PNP2 is the first PNP displaying a cytidine phosphorolysis activity in levels that might be relevant for cellular metabolism.

### *Sm*PNP2 crystal structures suggest a different gate loop dynamics in the active site

In order to obtain further insights in relation to *Sm*PNP2 nucleoside specificity, the crystallographic structure of the protein was obtained with different ligands. *Sm*PNP2 readily crystallized in several conditions, from pH 4.0 to pH 9.0, in cubic space group P2_1_3 with one monomer per asymmetric unit. Most of the *Sm*PNP2 crystals from co-crystallization experiments with cytidine show cytosine in the active site. Furthermore, a ternary complex with cytosine and R1P was also obtained, indicating the catalytic capability of *Sm*PNP2 in the crystallization solution and/or during incubation time. Soaking experiments using high ligand concentration was employed resulting in four new *Sm*PNP2 complexes with adenine, hypoxanthine, tubercidin, and cytidine. The data collection and refinement statistics are presented in Supplementary Table 1.

As expected, the overall fold of *Sm*PNP2 is the same observed for low molecular weight NPs, with a root-mean-square deviation (RMSD) of 1.17 Å when compared by superposition with trimeric *Sm*PNP1 structures (PDB IDs 1TCV, 3FAZ, and 3E0Q). The superposition of *Sm*PNP2 complex structures does not show any remarkable difference (RMSD of 0.47 Å for trimers and 0.33 Å for monomers). A larger RMSD was observed for *Sm*PNP2-tubercidin complex (0.82 and 1.24 Å for monomer and trimer, respectively). The main difference between the *Sm*PNP1 and *Sm*PNP2 structures is the presence of α-helix formed by residues 247-252 in the latter, at the beginning of the gate loop (residues 244-260).

Comparison between both *Sm*PNPs isoforms reveals an intriguing substitution, of a residue (A118L; *Sm*PNP1:*Sm*PNP2) belonging both to the Phosphate Binding Site (PBS) and a highly conserved motif of the NP-1 family members forming the “*bottom*” of the active site (13). This A118L substitution should have been preceded by a T244A substitution since a steric hindrance between side chains is predicted in a protein containing the T244 and an L118 (Figure 2). Erion et al (44), performed a site direct mutagenesis of T242A in human PNP, resulting in kinetic parameters for inosine and hypoxanthine similar to the wild-type enzyme, however, the K_M_ for phosphate is reduced 18 fold.

**Figure 2.**
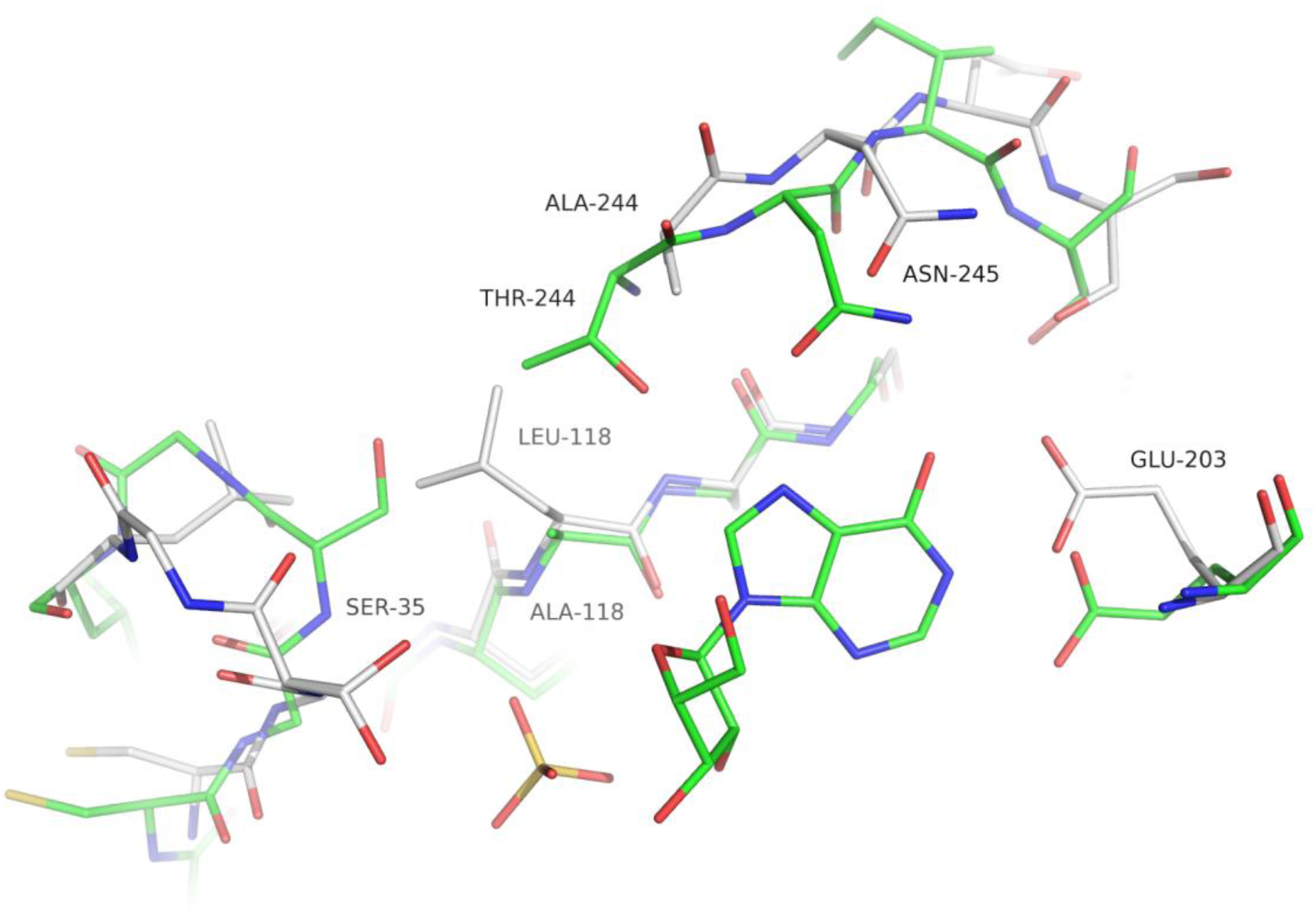
Comparison between *Sm*PNP1 (green backbone) and *Sm*PNP2 (white backbone) active site. The presence of A118L substitution results in the phosphate loop S35 displacement to open position due to steric hindrance and therefore the loop dynamic is no longer supported.

The presence of A118L substitution results in conformational changes of the phosphate loop centered in the S35 residue. This loop, corresponding to residues 32-40 in *Sm*PNP1, can assume two different conformations (closed and opened). The presence of a bulkier leucine group in place of alanine next to the S35 residue in *Sm*PNP2 block the movement necessary to assume a closed conformation due to steric hindrance maintaining this loop permanently in an open conformation (Figure 2).

Another relevant substitution in the active site region is the P257I at the ending of the gate loop. The ring of P257 restricts the conformations assumed by Y202 (which forms a pi-stacking within the base). The presence of I257 with displaces the residues Y202 and E203, and causes the reorientation of N245, reducing the volume of the active site cavity.

The presence of a new α-helix formed by residues 247-252 and substitution A118L also appear to have a large impact in the active site and gate loop conformations. This can be noted by calculating the RMSD of Cαs of residues 244-247, which display a values 2.84 Å (Figure 3A), thus indicating a considerable deviation from the structure of *Sm*PNP1. This displacement of residues 244-247 brings residue D250 in the vicinity of N245, resulting in an H-bond formation between N245 ND2 and D250 OD2, locking N245 in the new conformation (Figure 2 and 4). Consequently, the N245 side chain is unable to bind purine nucleosides in the same manner of *Sm*PNP1, as shown by the hypoxanthine, adenine (Figure 5), and tubercidin (Supplementary Figure 5) complex structures (discussed below). The residue N246 (I246 in *Sm*PNP1) appears to be determinant the appearance of the new α-helix in *Sm*PNP2, with the main chain N246 O and the OD1 atoms forming three hydrogen bonds (H-bonds) with main chain N of residues 248, 249 and 250 thus forming the N-cap of the α-helix, the ND2 of N246 also interacts with N123 O and L126 O (Figure 3B).

**Figure 3.**
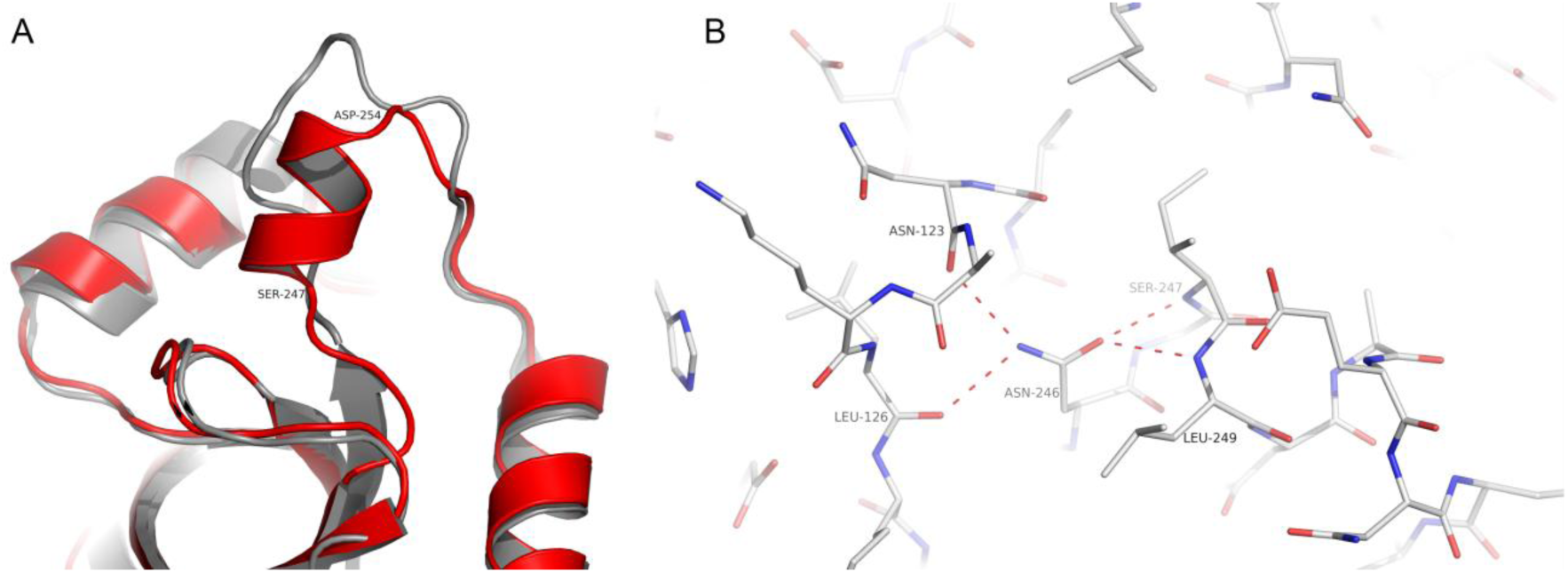
**A.** Ribbon model of *Sm*PNP1 (grey) and *Sm*PNP2 (red) superposition showing the extra α-helix formed by residues 247-252 in *Sm*PNP2. **B.** The formation of this α-helix is due to the presence of N246, which forms the N-terminus cap of the helix. The side chain of N246 forms 4 hydrogen bonds that lock this conformation.

**Figure 4.**
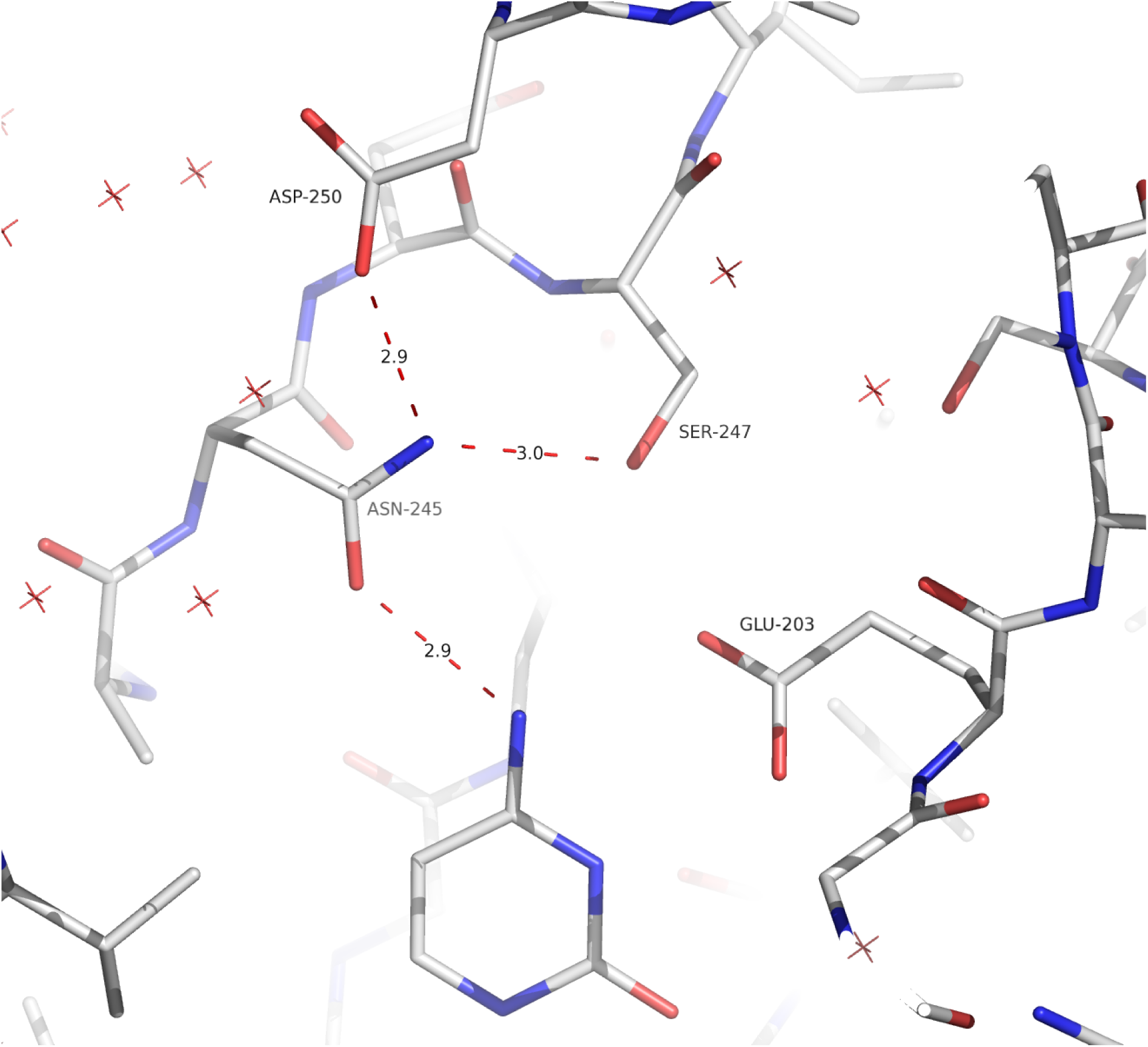
H-bond interaction between N245 and D250. This interaction helps N245 assuming this non-canonical conformation, rendering *Sm*PNP2 unable to bind purine nucleosides as observed for *Sm*PNP1. This interaction is possible due to the α-helix-formation by residue 246-252.

**Figure 5.**
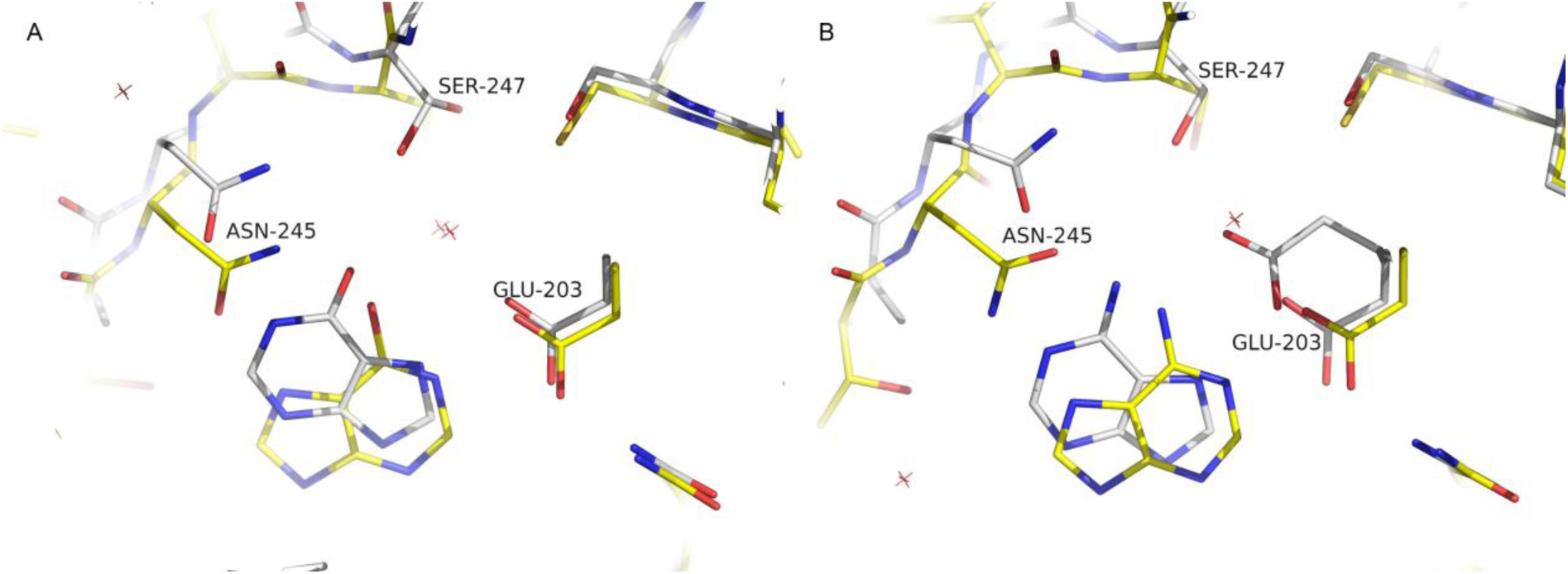
Purine bases in the *Sm*PNP2 active site show a very distinct orientation when compared to the *Sm*PNP1. **A.** Superposition of *Sm*PNP1 (yellow) and *Sm*PNP2 (white) hypoxanthine complexes. In *Sm*PNP2, hypoxanthine shows a completely different binding mode flipped 180° in comparison the canonical binding mode observed for *Sm*PNP1 and others LMM PNPs. The same scenario is observed for adenine. **B.** Superposition of *Sm*PNP1 (yellow) *Sm*PNP2 (white) adenine complexes. As observed for hypoxanthine, adenine binding mode resembles the first and also is flipped 180° in comparison to the *Sm*PNP1-adenine complex.

### Adenine and hypoxanthine bind *Sm*PNP2 active site in an unusual orientation

Using a soaking approach, three complexes were obtained: *Sm*PNP2-tubercidin (7-deaza-adenosine), *Sm*PNP2-hypoxanthine and *Sm*PNP2-adenine (The composite omit maps of these ligands can be seen in Supplementary Figure 6, 7A, and 7C, respectively). The interaction of *Sm*PNP2 with tubercidin in the ribose binding site (RBS) and PBS are similar to that observed for the *Sm*PNP1-adenosine complex. A notable exception is the distance of the base in respect to the E203 is greater than that observed in *Sm*PNP1 previous crystals (19), meaning that interaction between E203 and tubercidin in *Sm*PNP2 became water-mediated (E203 OE1 – w142-Tub N6) (Supplementary Figure 5A and 5B).

**Figure 6.**
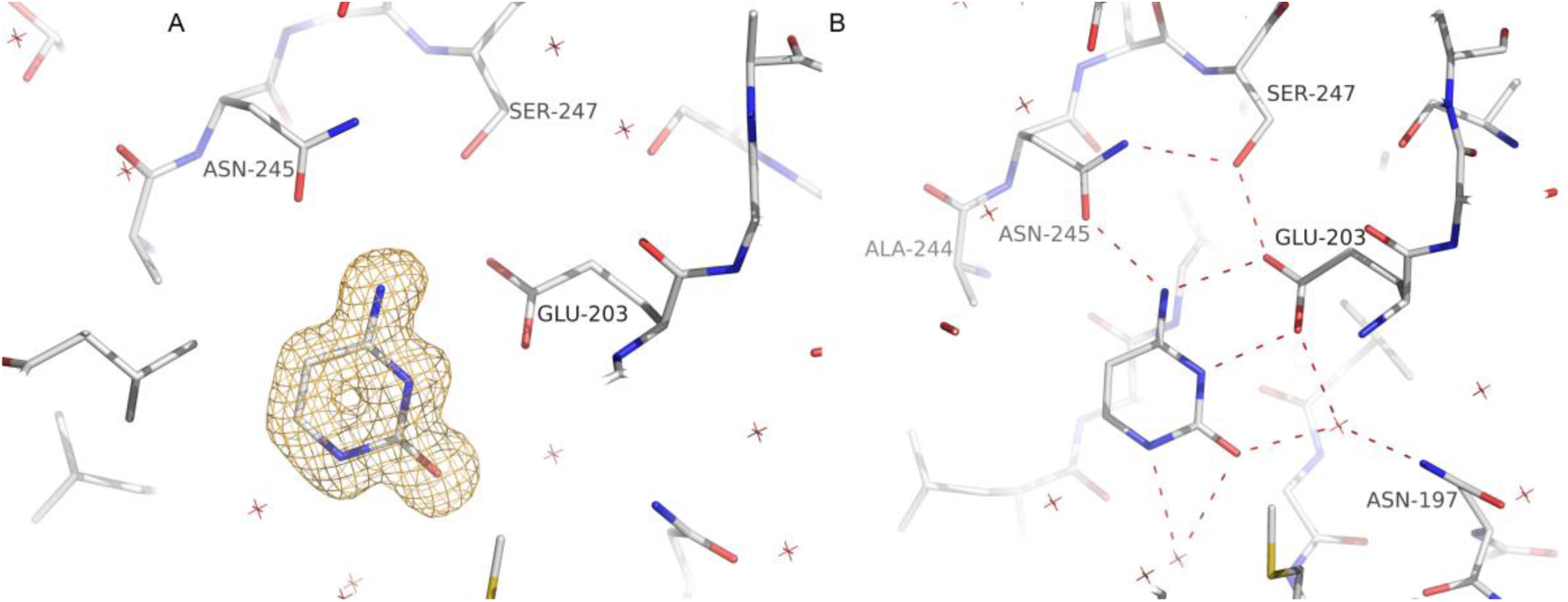
*Sm*PNP2 binds cytosine. **A.** Composite omit electron density map contoured at 1 σ for cytosine in the *Sm*PNP2 active site. **B.** Cytosine H-bond interactions network in the *Sm*PNP2 active site. The cytosine molecule forms 6 H-bonds with active site residues and water molecules (represented in red X).

**Figure 7.**
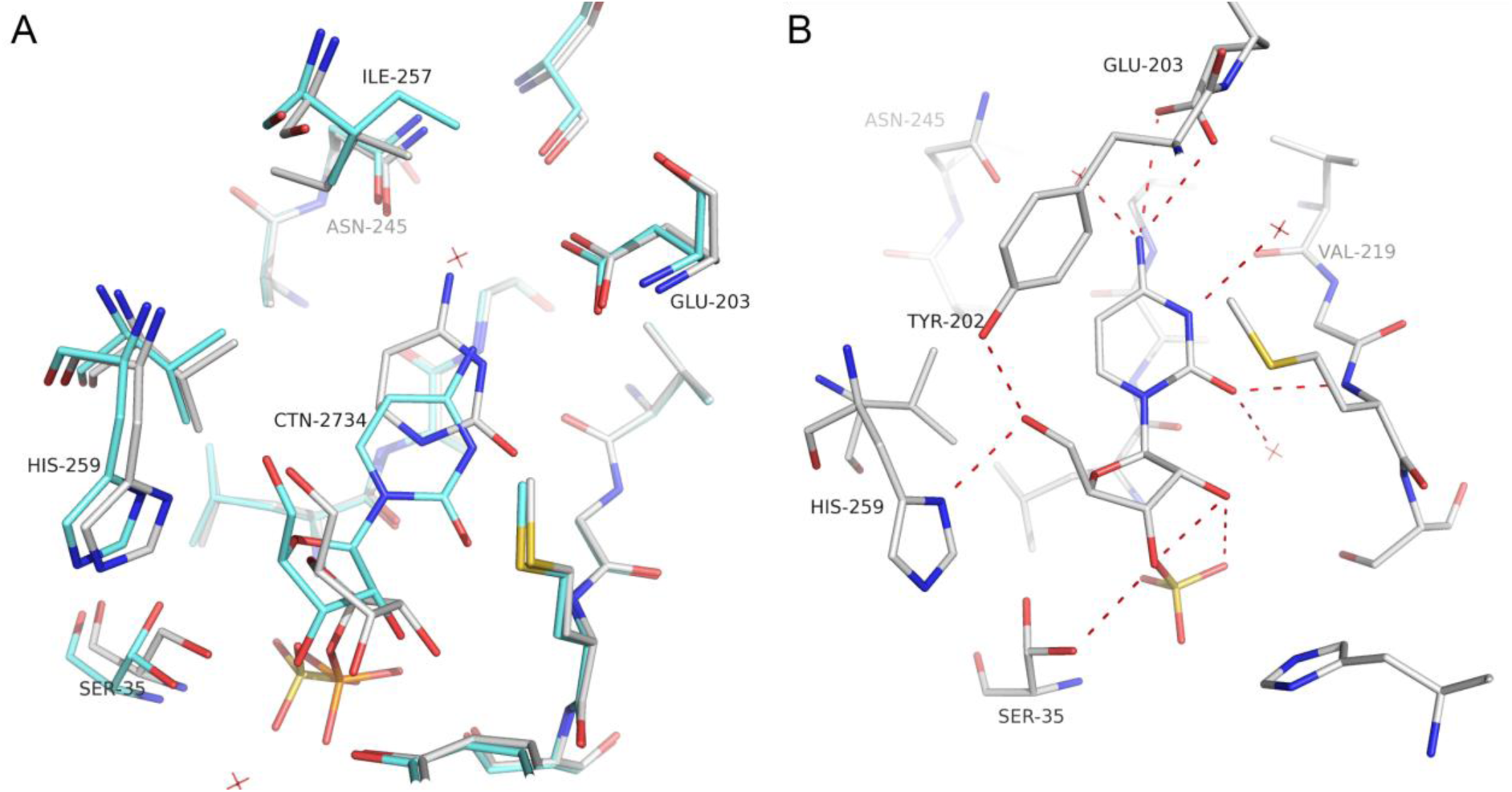
**A.** Superposition of cytidine (blue) and cytosine-R1P (white) *Sm*PNP2 complexes. **B.** Cytidine H-bond interactions in the *Sm*PNP2 active site. Differently, of other nucleosides, no direct interaction was formed between the base and N245, the interaction is now mediated by a water molecule (represented in red X).

A totally different scenario is observed for the binding of adenine or hypoxanthine in the *Sm*PNP2, where the bound base show a 180° rotation in Y axes followed a small rotation in the Z axes resulting in different interactions within BBS in comparison to PNP1 (Figure 5). A water molecule that interacts with E203, S247 and the base in *Sm*PNP1 was also observed in the *Sm*PNP2-hypoxanthine complex (Supplementary Figure 7B). The binding of adenine is similar to the hypoxanthine, however, a clear double conformation is observed for E203 (one of them is the canonical one), in this configuration the side chain of E203 interacts only with N7 (canonical conformation) (Supplementary Figure 7D).

Considering that tubercidin, which is an adenosine analog, display a canonical binding mode, it is possible to speculate that both adenosine and inosine should be also binding in a canonical position. The main reason for that is the sterical constraints related to the presence of ribose group, which would argue against a similar conformation seen for adenine and hypoxanthine. This would mean that only after the glycoside bond cleavage the base could be free to rotate in the BBS and assume a non-canonical orientation.

### *SmPNP*2 crystal structures confirm specific ligation of cytosine and cytidine to the active site

Analysis of the *Sm*PNP2-cytosine complex structure obtained at 1.36 Å places the cytosine molecule in the active site (Figure 6A), where it forms 6 H-bonds. The side chain of key active site residues E203 and N245 forms two and one H-bonds, respectively, and other three water-mediated bonds were found with residues L118, N197, and M221 (Figure 6B). The conformation of the BBS is the same observed for *Sm*PNP2 apo structure, and both structures are essentially the same.

The ternary *Sm*PNP2 – cytosine – R1P complex was also obtained at high resolution (1.42 Å) and it is well superimposed to *Sm*PNP2 – cytosine structure with RMSD of 0.19 Å. Both cytosine and R1P does not present a sharp electron density map as expected for this resolution, indicating movement of these ligands in the active site and/or the presence of cytidine and cytosine/R1P at the same time in the crystal. This is true especially for the ribose moiety of the R1P (Supplementary Figure 8).

The binding of cytosine and R1P causes small rearrangements in the active site when compared with the *Sm*PNP2-cytosine complex. A small displacement of E203 side chain occurs, increasing the H-bond distance (0.3 and 0.4 Å increase) with the cytosine resulting in a weak interaction. The interaction with N245 is maintained unaffected. The R1P binds in *Sm*PNP2 is a similar way as observed for *Sm*PNP1-R1P complex (24) as expected.

A more noteworthy modification in the binding site is observed for the *Sm*PNP2-cytidine complex. Comparison with both *Sm*PNP2 cytosine and cytosine and R1P complexes reveals that RMSD between cytosine moieties in both complexes is 1.73 Å (Figure 7A).

Interaction between *Sm*PNP2 and the cytosine moiety in cytidine occur through two direct H-bonds with residues M221 and E203 and three water-mediated anchored by residues E203 and N245 (water 3), E203 and N197 (water 29), and M221, L118, Y194 and S222 (water 45), this later also interacts with sulphate group which lies in phosphate binding site (Figure 7B).

The position of the ribose moiety in *Sm*PNP2-cytidine complex is displaced in comparison to what is observed in other *Sm*PNP1 and *Sm*PNP2 structures. Consequently, several side groups are moved to accommodate this new position from ribose. The H259 ND1 and Y202 OH are both displaced when compared with the *Sm*PNP2-cytosine-R1P to form H-bonds to ribose O5. Small movements are also observed for the phosphate S35 loop and for H88 side chain. Moreover, a structural water that typically mediates the interaction between Y194 and ribose O2’ interacts with cytidine O2 in *Sm*PNP2-cytidine complex and the canonical contacts with M221 and Y90 are lost in this complex (Figure 7B).

## Discussion

The unique properties of *Sm*PNP2 were discovered using a combination of techniques as DSF, kinetics and X-ray crystallography. It is noteworthy that the ability of *Sm*PNP2 to catalyze the phosphorolysis of cytidine with similar efficiency as inosine, which represents a unique trait in a nucleoside phosphorylase. The crystal structure reveals this property appears to be related to the substitutions in the active site that induce the locking of the gate loop. Indeed, the mobility of active site loops has been ascribed to play a key role in mediating enzyme promiscuity (45), providing further support to this hypothesis.

Crystal structures of *Sm*PNP2 may also help to contribute to the understanding of catalysis in PNPs since different and sometimes contrasting catalytic mechanism has been proposed for phosphorolysis in LMM NPs (46). For example, mechanisms that involve the protonation or stabilization of N(7) from the purine ring promoted by N245 (N243, in human) (47, 48) are not expected to occur within the structural framework observed in *Sm*PNP2, since this residue is now locked in new conformation and therefore a H-bond between N245 side chain and N(7) is absent.

The fact that *Sm*PNP2 is capable of hydrolyzing adenosine and inosine with similar efficiency also exclude a mechanism proposed for HMM PNPs based on transition state where E204 (E201 in human PNP) forms an H-bond with exocyclic negatively charged O(6) of rare enol purine tautomer (49). This is because in such model adenosine could not be used as a substrate since no stabilization is possible with a negative charge at the base via enolate intermediate and the lack of hydrogen at N1 of the base.

In contrast, an alternative mechanism proposed for HMM PNPs based on the formation of a negative charge delocalized to the base six-membered ring during transition state and a network of water molecules that connect N1 and O6 of the purine base (50), seems compatible with *Sm*PNP2 traits. That is because that mechanism allows for catalysis of both 6-oxo and 6-amino purines and *Sm*PNP2 display a water molecule anchored by residues E203 and S247, which would help in the stabilization of a charged intermediate, together with E201 side chain.

Moreover, a proposed catalytic mechanism for *Trypanosoma cruzi* uridine phosphorylase (51) seems an adequate mechanism to describe the mechanism for cytidine cleavage by *Sm*PNP2. Under the original model a protonated group, possibly uridine 3-NH, is essential for catalysis, due their deprotonation generates a dianionic uracil a less effective leaving group. In the context of the catalysis promoted by *Sm*PNP2, we can assume that an analogous configuration could occur with E203 forming 2 H-bond with cytidine N4 and one water molecule also binding N4 anchored by residues E201 and N245. In this context, a protonation of N3 could achieve helped by a water molecule anchored by residue N197.

Analysis of Lu and co-workers (52, 53) data related to gonad-specific and pairing-dependent transcriptome reveal that *Sm*PNP1 is approximately 160 times more expressed than *Sm*PNP2 in mature adult females. This is probably related to the fact that *Sm*PNP1 display high expression in female vitelaria, as revealed by our WISH experiments. Moreover, *Sm*PNP2 is significantly enriched in male’s testis when compared to its own expression in whole adult males (~25 times higher expression). This data suggest that both PNPs might have an important role in functions related with sexual tissue metabolism.

A possible explanation of *Sm*PNP relative promiscuity in relation to the bound nucleotide would be a function related to scavenging of nutrients (15). Cytidine is the pyrimidine nucleoside with the highest incorporation in adult worms (210 pM per 10 worms pairs) (11) and is used by the uridine-cytidine kinase to form CMP in the worm metabolism. In cercariae, where *Sm*PNP2 has the highest transcript levels, this enzyme could be involved in the production of R1P from nucleosides that could be subsequently used by to form PRPP in order to save nucleosides.

These structures and kinetics of *Sm*PNP2 are another layer of information about the nucleotide metabolism in general and in particular for *S. mansoni* which enrich the understanding of the utilization of nucleotides for this important neglected parasite.

## Acknowledgments

We acknowledge the Fundação de Amparo a Pesquisa do Estado de São Paulo (FAPESP) grants 2012/10213-9 (JRT), 2012/05532-8 (LR), 2012/23730-1 (VHBS), 2012/14223-9 (HMP) and CNPq grant 474402/2013-4 for financial support. We also acknowledge Leticia Anderson and Sergio Verjovski-Almeida for providing adult *S. mansoni* worms to support the WISH experiments.

## Supplementary table

**Table 1.** Data collection, processing and refinement parameters.

|  | *SmPNP2-apo* | *SmPNP2-MES* | *SmPNP2-CYT* | *SmPNP2-CYT-R1P* |
| --- | --- | --- | --- | --- |
| Detector | Pilatus 2M | Pilatus 2M | Pilatus 6M | Pilatus 6M |
| Cell parameters (Å) a; b; c | 99.55; 99.55; 99.55 | 97.78; 97.78; 97.78 | 99.13; 99.13; 99.13 | 98.64; 98.64; 98.64 |
| Space Group | P2<sub>1</sub>3 | P 2<sub>1</sub>3 | P 2<sub>1</sub>3 | P 2<sub>1</sub>3 |
| Resolution (Å) | 44.52 - 1.57 (1.61 - 1.57) | 43.73 - 1.74 (1.79 - 1.74) | 70.10 - 1.36 (1.41 - 1.36) | 69.75-1.42 (1.46 - 1.42) |
| X-ray Source | DLS I04-1 | DLS I04-1 | DLS I04 | DLS I04 |
| $\lambda$ (Å) | 0.920 | 0.920 | 0.979 | 0.979 |
| Multiplicity | 6.8 (7.0) | 11.0 (10.5) | 10.6 (10.0) | 11.4 (9.1) |
| R<sub>merge</sub> (%) | 5.5 (62.3) | 5.8 (68.9) | 13 (93.1) | 7.6 (147.3) |
| R<sub>pim</sub> (%) | 2.5 (27.1) | 1.9 (23.3) | 4.2 (46.2) | 2.4 (56.0) |
| CC(1/2) | 0.998 (0.832) | 1 (0.844) | 0.994 (0.60) | 0.999 (0.540) |
| Completeness (%) | 100 (100) | 100 (100) | 100.0 (100.0) | 100 (99.9) |
| Reflections | 312485 (23781) | 354427 (22974) | 737563 (43532) | 689682 (40358) |
| Unique Reflections | 46089 (3379) | 32239 (2370) | 69723 (7274) | 60417 (4438) |
| I/$\sigma$ | 18.0 (2.9) | 25.4 (3.6) | 8.4 (1.5) | 15.9 (1.3) |
| Reflections used for Refinement | 46050 | 31691 | 69526 | 60379 |
| R(%)** | 18.53 | 15.0 | 17.46 | 16.19 |
| R<sub>free</sub>(%)** | 21.24 | 19.11 | 19.16 | 17.85 |
| N° of protein atoms | 2180 | 2173 | 2541 | 2550 |
| N° of ligant atoms | - | 11 | 29 | 30 |
| B (Å<sup>2</sup>) | 18.79 | 17.33 | 15.64 | 17.93 |
| Coordinate Error (ML based) (Å) | 0.15 | 0.18 | 0.18 | 0.16 |
| Phase error (°) | 21.18 | 17.07 | 21.68 | 17.39 |
| *Ramachandran Plot* |  |  |  |  |
| Favored (%) | 97.54 | 98.94 | 98.6 | 97.89 |
| Allowed (%) | 2.11 | 1.06 | 1.40 | 2.11 |
| Outliers (%) | 0.35 | 0.0 | 0.0 | 0.0 |
| All-atom Clashscore | 1.81 | 3.38 | 4.27 | 2.92 |
| *RMSD from ideal geometry* |  |  |  |  |
| r.m.s. bond lengths (Å) | 0.007 | 0.018 | 0.005 | 0.006 |
| r.m.s. bond angles (°) | 1.088 | 1.694 | 0.872 | 0.953 |
| *PDB ID* | 5CXQ | 5CXS | 5KO5 | 5KO6 |

|  | *Sm*PNP2-Adenine | *Sm*PNP2-Cytidine | *Sm*PNP2-Hypoxanthine | *Sm*PNP2-Tubercidin |
| --- | --- | --- | --- | --- |
| Detector | Pilatus 6M | Pilatus 6M | Pilatus 6M | Pilatus 6M |
| Cell parameters (Å) a; b; c | 100.83; 100.83; 100.83 | 100.14; 100.14; 100.14 | 99.70; 99.70; 99.70 | 91.98; 91.98; 91.98 |
| Space Group | P2<sub>1</sub>3 | P 2<sub>1</sub>3 | P 2<sub>1</sub>3 | P 2<sub>1</sub>3 |
| Resolution (Å) | 71.30 – 1.90 (1.94 – 1.90) | 57.82 – 2.10 (2.16 – 2.10) | 70.00 – 2.10 (2.16 – 2.10) | 65.04 – 1.95 (2.00 – 1.95) |
| X-ray Source | DLS I04-1 | DLS I04-1 | DLS I04-1 | DLS I04-1 |
| $\lambda$ (Å) | 0.920 | 0.920 | 0.920 | 0.920 |
| Multiplicity | 7.6 (7.8) | 6.9 (7.0) | 6.7 (4.7) | 5.4 (5.3) |
| R<sub>merge</sub> (%) | 7.0 (77.9) | 18.2 (140.1) | 15.4 (75.4) | 8.7 (74.1) |
| R<sub>pim</sub> (%) | 2.9 (32.1) | 7.9 (61.5) | 6.9 (42.4) | 4.5 (39.2) |
| CC(1/2) | 1.000 (0.803) | 0.994 (0.598) | 0.993 (0.627) | 0.998 (0.710) |
| Completeness (%) | 100.0 (100.0) | 99.9 (100.0) | 99.9 (99.1) | 100.0 (100.0) |
| Reflections | 205704 (13435) | 137829 (11377) | 130953 (7589) | 104475 (7301) |
| Unique Reflections | 27201 (1720) | 19849 (1614) | 19582 (1610) | 19223 (1368) |
| I/$\sigma$ | 18.7 (2.6) | 7.2 (1.5) | 9.1 (1.8) | 9.4 (1.8) |
| Reflections used for Refinement | 27154 | 19795 | 19553 | 19152 |
| R(%)** | 16.00 | 19.04 | 18.87 | 17.87 |
| R<sub>free</sub>(%)** | 19.68 | 22.60 | 22.75 | 20.24 |
| N° of protein atoms | 2174 | 2170 | 2170 | 2158 |
| N° of ligand atoms | 25 | 23 | 22 | 28 |
| B (Å<sup>2</sup>) | 26.01 | 20.00 | 20.92 | 28.20 |
| Coordinate Error (ML based) (Å) | 0.22 | 0.22 | 0.25 | 0.18 |
| Phase error (°) | 18.55 | 22.93 | 21.49 | 22.86 |
| *Ramachandran Plot* |  |  |  |  |
| Favored (%) | 98.59 | 98.94 | 98.59 | 98.58 |
| Allowed (%) | 1.06 | 1.06 | 1.41 | 1.42 |
| Outliers (%) | 0.35 | 0.00 | 0.00 | 0.00 |
| All-atom Clashscore | 2.71 | 2.26 | 1.35 | 1.81 |
| *RMSD from ideal geometry* |  |  |  |  |
| r.m.s. bond lengths (Å) | 1.205 | 0.604 | 0.002 | 0.010 |
| r.m.s. bond angles (°) | 0.015 | 0.002 | 0.524 | 1.068 |
| *PDB ID* | 5TBS | 5TBT | 5TBU | 5TBV |

## Supplementary Figures

**Supplementary figure 1.**
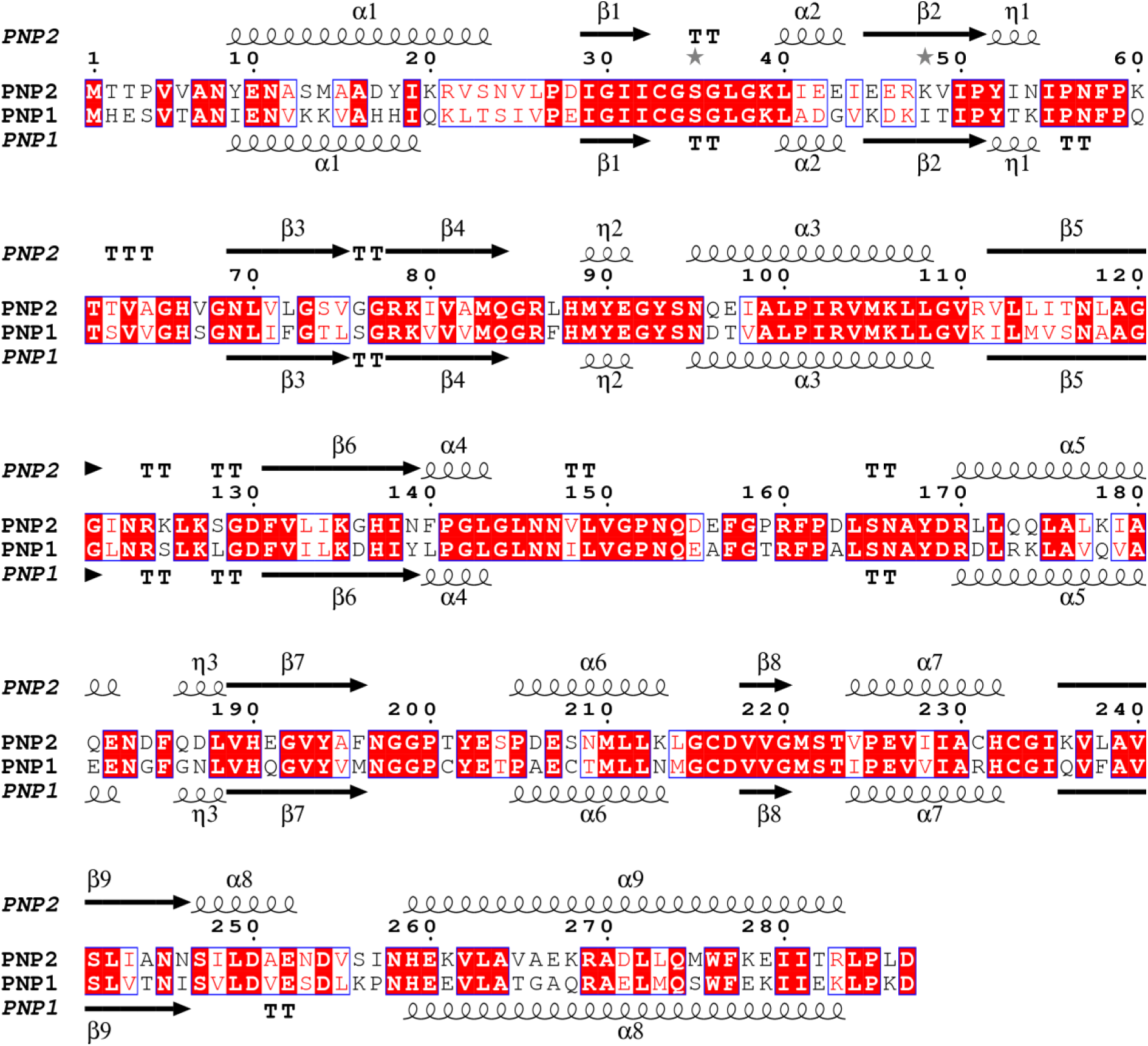
Alignment of *Sm*PNP1 and *Sm*PNP2 sequences, where is important to highlight the presence of an extra α-helix at residue 250 in *Sm*PNP2 structure. The sequences share 61% identity.

**Supplementary figure 2.**
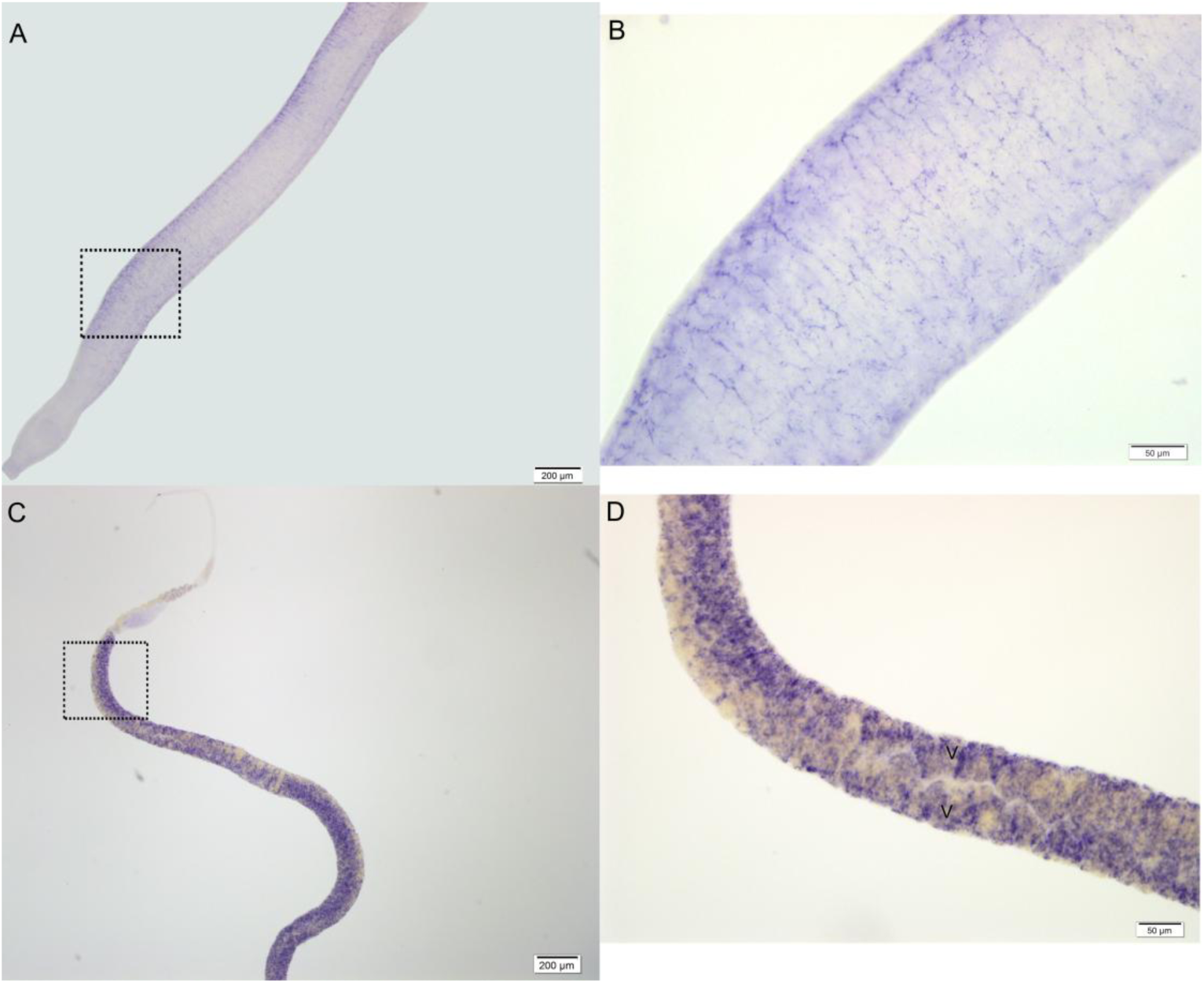
Localization of PNP1 transcripts in *S. mansoni* adult worms by WISH. PNP1 expression sites in male (A-B) and female (C-D) adult worms. B and D are higher magnification views of the boxed images in A and C, respectively. V, vitellaria of female worms.

**Supplementary figure 3.**
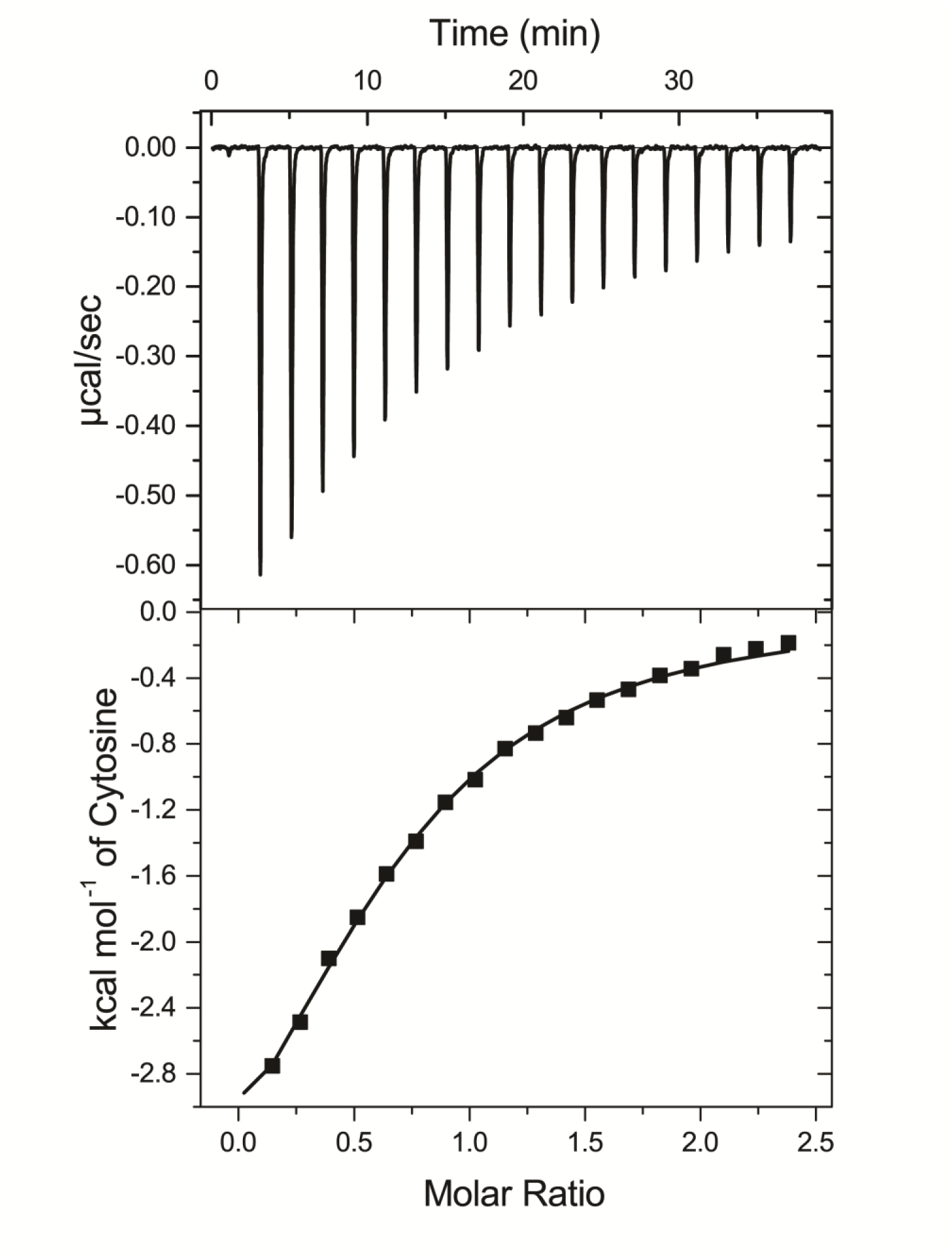
ITC binding curve of *Sm*PNP2 and cytosine base. Top panel: Curve of the titration of 60 μM of *Sm*PNP2 with cytosine 1 mM at 25 °C. Thermodynamic parameters were derived from non-linear least-squares fitting. Bottom Panel: fit of the binding isotherm to the one set of sites model.

**Supplementary figure 4.**
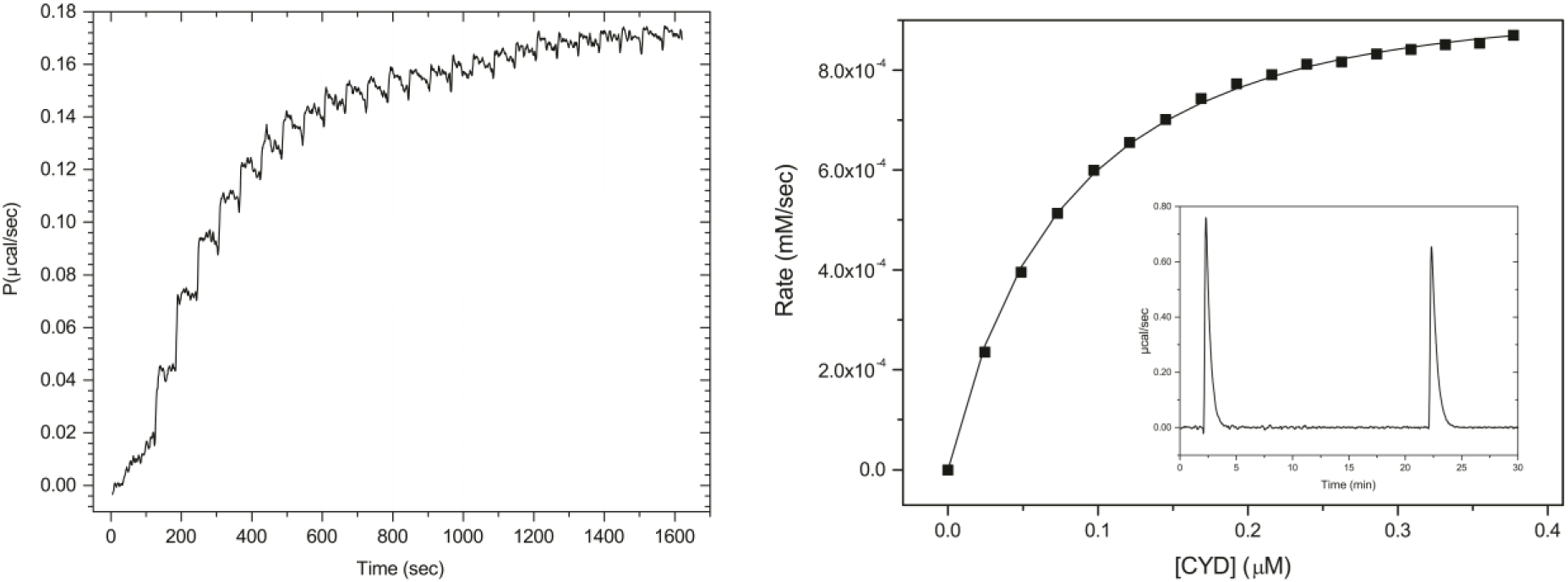
**A.** Multiple-injection titration of *Sm*PNP2 13 μM with cytosine 5 mM; **B.** Fit of the Michaelis–Menten model (solid line) to the *Sm*PNP2 protein reaction rate as a function of the added cytosine derived from the ITC data. *Inset*: Apparent enthalpy change experiment determination for the catalytic reaction.

**Supplementary figure 5.**
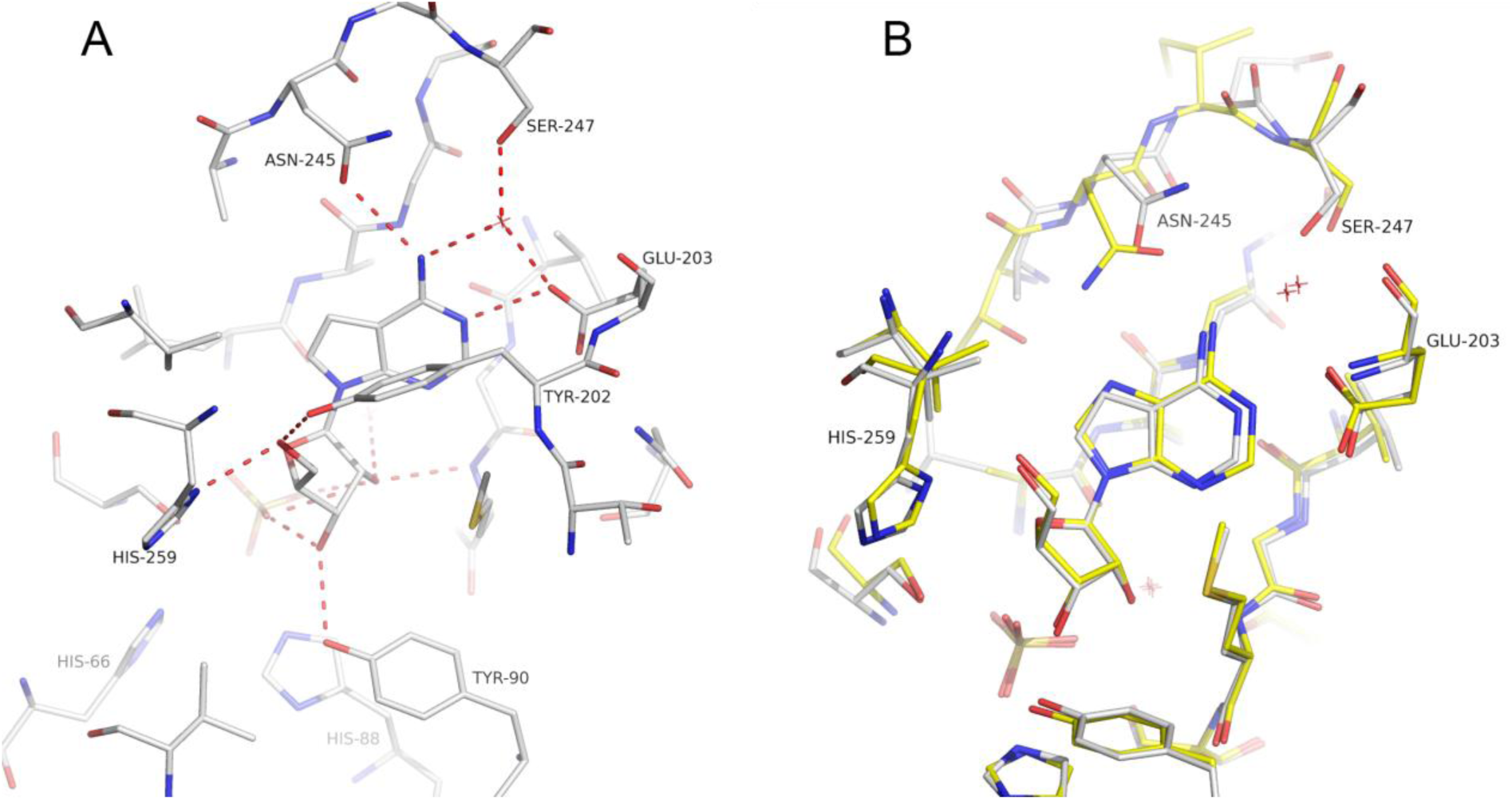
Tubercidin binding in *Sm*PNP2 active site. **A**. Tubercidin interactions in *Sm*PNP2 active site. Different to adenosine binding in *Sm*PNP1, no direct interaction was formed between E203 side chain and tubercidin. **B**. Superposition of *Sm*PNP1-adenosine (yellow) and *Sm*PNP2-tubercidin (white), with exception of different conformers for N245 side chain no other differences were observed, in this complex E203 assumes the canonical conformation.

**Supplementary figure 6.**
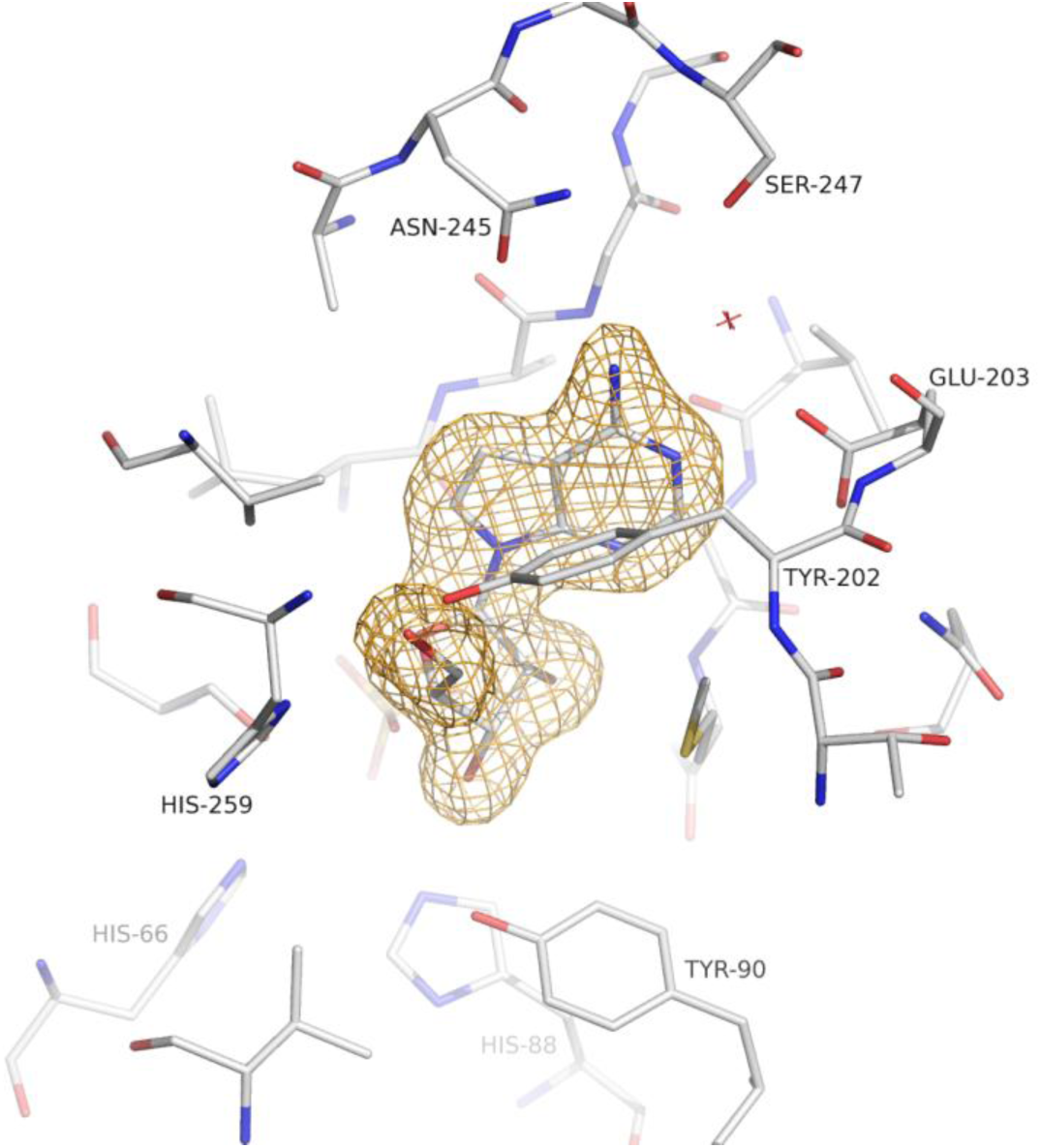
Composite omit map countered at 1.0 σ for tubercidin in *Sm*PNP2 active site.

**Supplementary figure 7.**
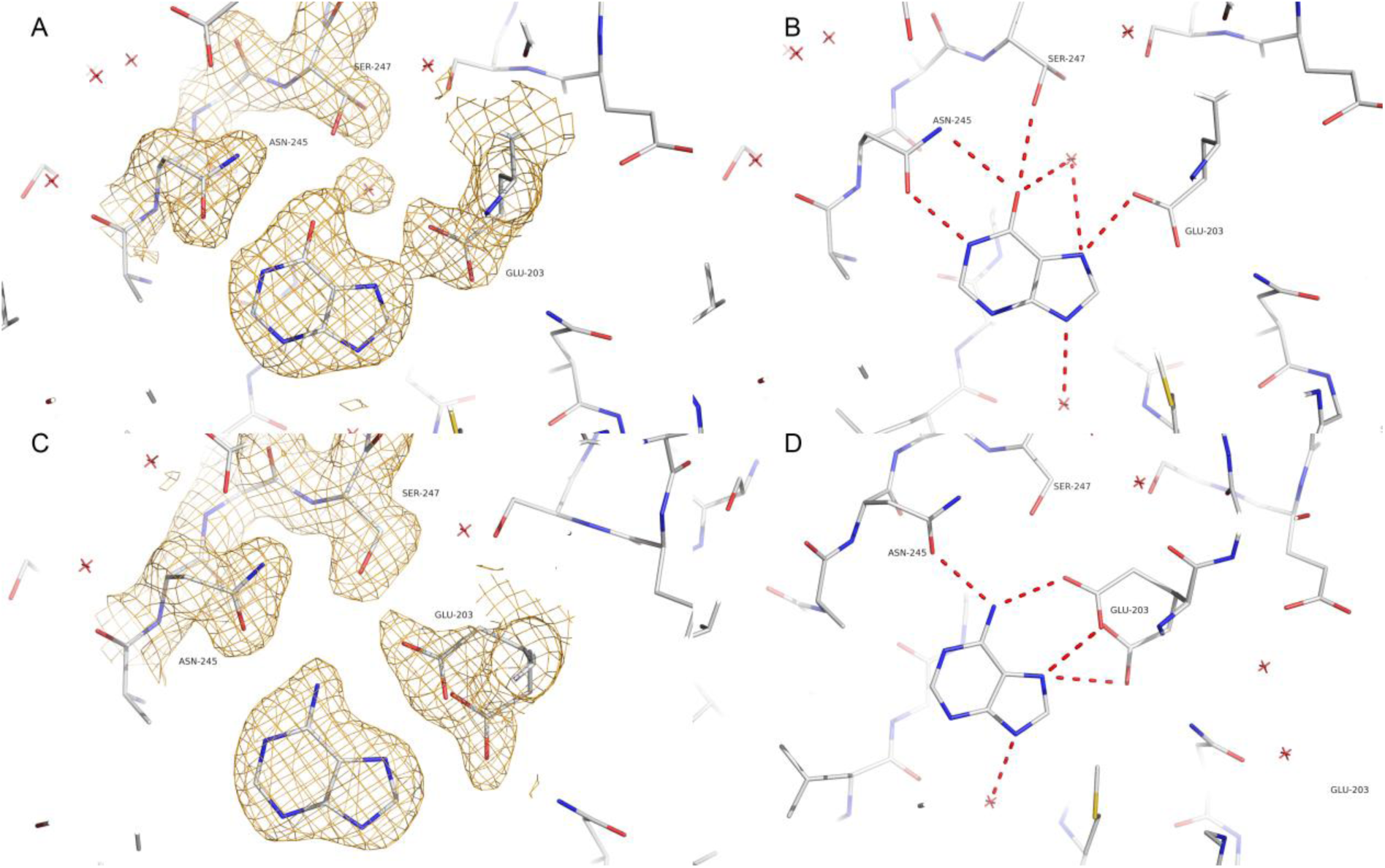
**A.** Composite omit map countered at 1.0 σ for *Sm*PNP2 – hypoxanthine complex. **B**. Hypoxanthine H-bond interaction formed in the *Sm*PNP2 active site. **C.** Composite omit map countered at 1.0 σ for adenine. **D.** Adenine H-bond interactions in the *Sm*PNP2 active site.

**Supplementary figure 8.**
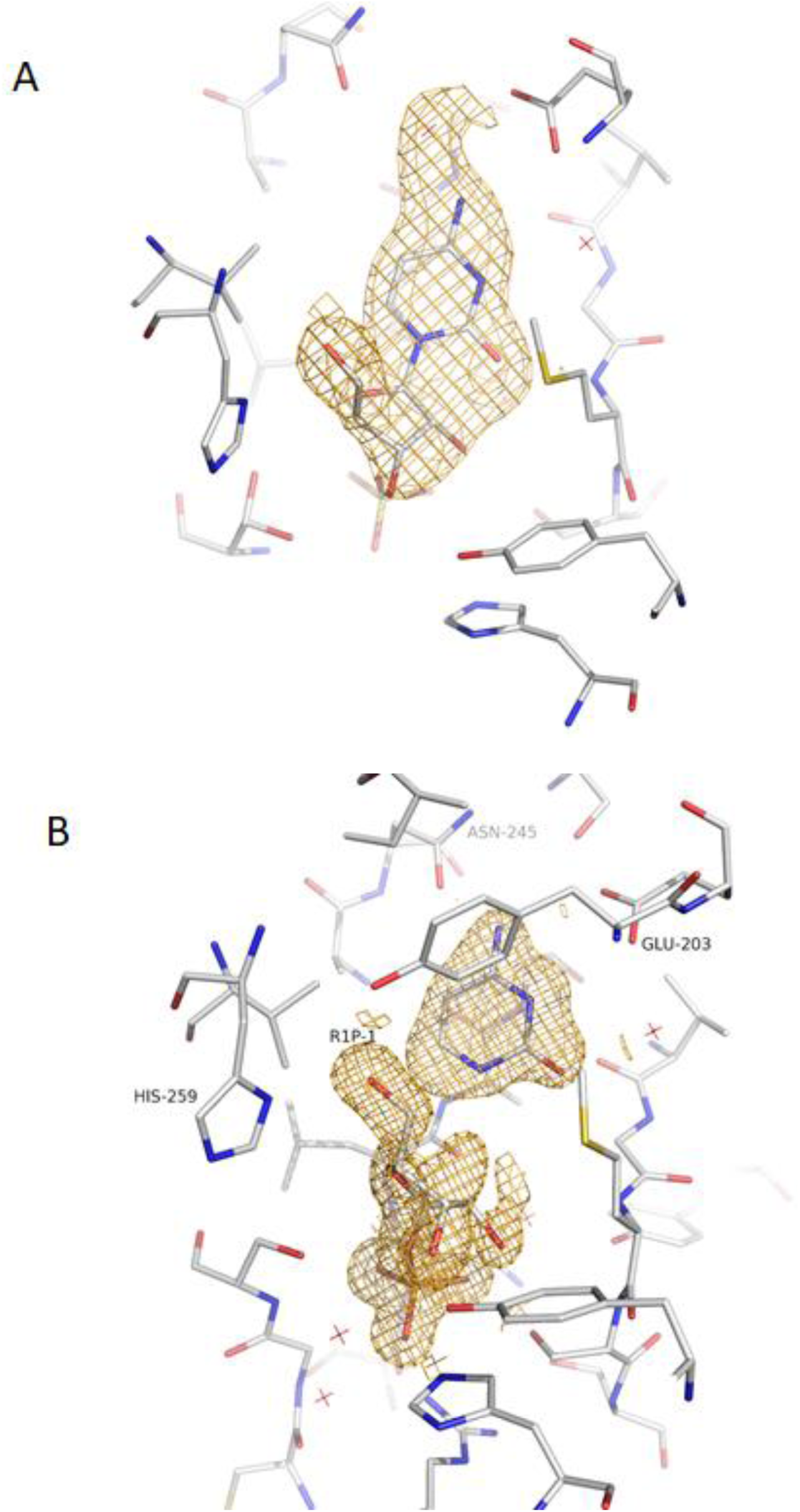
**A.** Composite omit map countered at 1.0 σ for *Sm*PNP2 – cytidine. **B.** Composite omit map countered at 1.0 σ for *Sm*PNP2 – ribose-1-phosphate and cytosine complex.

